# Empirical characterization factors assessing the effects of hydroelectricity on fish richness across three large biomes

**DOI:** 10.1101/678383

**Authors:** Katrine Turgeon, Gabrielle Trottier, Christian Turpin, Cécile Bulle, Manuele Margni

## Abstract

Hydroelectricity is often presented as a clean, reliable, and renewable energy source, but is also recognized for its potential impacts on aquatic ecosystem biodiversity. We used empirical data on change in fish species richness following impoundment to develop Characterisation Factors (CF) and Impact Scores (IS) for hydroelectricity production for use in Life Cycle Assessment (LCA). We used data collected on 89 sampling stations (63 upstream and 26 downstream of a dam) belonging to 27 reservoirs from three biomes (boreal, temperate and tropical). Overall, the impact of hydroelectricity production on fish species richness was significant in the tropics, of smaller amplitude in temperate and minimal in boreal biome, stressing for the need of regionalisation. The impact of hydroelectricity production was also quite consistent across scales (*i.e.*, same directionality and statistical significance across sampling stations, reservoirs and biomes) but was sensitive to the duration of the study (*i.e.*, the period over which data have been collected after impoundment), highlighting the need for a clear understanding of transient situations before reaching steady states. Our CFs and ISs contribute to fill a gap to assist decision makers using LCA to evaluate alternative technologies, such as hydropower, to decarbonize the worldwide economy.

**Highlights:**

- This paper is the first to develop global and empirically based characterization factors of the impact of hydroelectricity production on aquatic ecosystems biodiversity, to be used in LCA;
- The impact of hydroelectricity production on fish species richness was significant in the tropics, of smaller amplitude in temperate and minimal in boreal biome;
- The impact of hydroelectricity production on fish richness was consistent across scales - same directionality and statistical significance across sampling stations, reservoirs and biomes;
- The impact of hydroelectricity production on fish richness was sensitive to the duration of the study, highlighting the need for a clear understanding of transient situations before reaching steady states in LCA.

## 1. Introduction

One of the most important challenge we face as a society is the increased demand for energy (SEforALL, 2016, p. 4). In response to this worldwide demand, hydroelectricity is presented as a relatively clean, reliable, and renewable energy source (Tahseen and Karney, 2017; Teodoru et al., 2012), and an interesting option to decarbonise our global economy (Figueres et al., 2017; Potvin et al., 2017) by reducing greenhouse gas emissions (GHGs). Hydroelectricity supplies less than 3% of the primary energy worldwide but more than 70% of the world’s renewable electricity (International Energy Agency, 2017; World Energy Council, 2016). These numbers will increase in the coming years as many large dams are being constructed around the world, particularly in developing economies that are mostly located in the tropics (Grill et al., 2015; Winemiller et al., 2016).

Despite its recognized advantages, the production of hydroelectricity can impact aquatic ecosystem functions and biodiversity through the regulation of the river flow, by a drastic changes in the hydrological regime, and by the fragmentation of rivers (Gracey and Verones, 2016; Renöfalt et al., 2010; Rosenberg et al., 2000). Dams constructed for hydroelectricity production, transform large rivers (*i.e.*, lotic environment) and surrounding lakes into larges reservoirs (*i.e.*, lentic environment), or a series of reservoirs (*sensu* cascade reservoirs; (Friedl and Wüest, 2002; Haxton and Findlay, 2009). Upstream of the dam, reservoirs can experience variation in water levels outside of their natural amplitudes (Kroger, 1973; Zohary and Ostrovsky, 2011). Downstream of the dam, changes in seasonal and inter-annual streamflow magnitude and variability are generally reduced (Friedman et al., 1998; Graf, 2006) and fish movement can be altered by the dam. These modifications can impact the biodiversity, abundance, distribution and community structure of many taxa of the aquatic food web (Furey et al., 2006; Nilsson and Berggren, 2000; Vörösmarty et al., 2010).

Life cycle assessment (LCA) is used to assess the environmental impacts of products and services throughout their whole life cycle (*i.e.*, cradle-to-grave; Finnveden et al., 2009; ISO, 2006). LCA informs about environmentally sound choices in the context of decision-making and is based on scientific evidence. When compared to other electricity production technologies, hydroelectricity scored favorably in LCA studies regarding GHG emissions, air pollution, health risk, acidification and eutrophication of ecosystems (CIRAIG, 2014; Hertwich, 2013; Sathaye et al., 2011). However, some of the impacts of hydroelectricity production on ecosystems and biodiversity are still not successfully integrated into LCA and are underrepresented due to some methodological challenges (de Baan et al., 2013; Gracey and Verones, 2016).

Evaluating and including the impacts of hydroelectricity production on aquatic ecosystems quality in LCA has been proven to be challenging because of the large data requirement, unclear causational effects, incomplete coverage of biodiversity impacts, and spatial and temporal scaling issues that can hinder its global application and validity (Gracey and Verones, 2016; McManamay et al., 2015; Milà i Canals et al., 2009; Teixeira et al., 2016). Different indicators have been proposed to measure the impacts on ecosystems and biodiversity in LCA (*e.g.*, difference in species richness, *i.e.*, the number of species, ecosystem scarcity and vulnerability, functional diversity; (Curran et al., 2011; Souza et al., 2013). But experts concluded – without a clear consensus – that change in species richness is a good and simple starting point to assess biodiversity impacts (Teixeira et al., 2016).

When change in species richness is used in LCA, it is essential to adequately consider the right spatial and temporal scale of impacts. Patterns observed locally (*e.g.*, in a reservoir) cannot always be extrapolated within or across regions. It is also important to evaluate the impact at the steady state, *i.e.*, at the time at which change in biodiversity stabilize after impoundment. Very few studies examined global impacts of hydroelectricity on ecosystems quality, or examined if patterns can be extrapolated across scales (but see (de Baan et al., 2013) for a multiple spatial scale study), and no study yet use empirically derived Characterization Factor (CF) and Impact Score (IS).

Here, we used empirically derived rate of change in fish species richness over time, across 89 sampling stations, belonging to 27 storage reservoirs from boreal, temperate and tropical biomes. The focus of this study is on storage reservoirs because of a lack of adequate longitudinal data (data before and after damming) from the other technologies (*e.g.*, run of the river and pumping stations). Our goals were to: 1) develop robust empirical CFs across three spatial scales (sampling station, reservoir and biome), 2) calculate the impact score of the creation of a reservoir (ISR) and of hydroelectricity production (IS) across scales, and 3) to test the need for regionalisation by examining if the observed patterns were consistent across biomes.

## 2. Materials and Methods

### 2.1 General approach

The approach to generate Characterization factors (CF) and Impact Scores (IS) was based on the examination of empirical patterns of changes in fish richness in response to river impoundment across three large biomes (boreal, temperate and tropical) from an extensive literature search. To calculate CF, we used the Potentially Disappeared Fraction of species (PDF) as the unit to express change in richness in response to hydroelectricity production. This unit has the advantage to be compatible with other damage oriented impact assessment methods addressing ecosystems quality such as IMPACT 2002+ (Jolliet et al., 2003) and Impact World + (Bulle et al., 2019) has been recommended by the UNEP-SETAC Life Cycle Initiative as an adequate and consistent biodiversity attribute (Verones et al., 2017). Because no reliable data were available to evaluate the biodiversity recovery when powerplants are decommissioned and dam removed, we were not able to address the recovery phase in the LCA and therefore focused our effort on the impacts during the occupation phase (*i.e.*, time span covering the construction of the dam until complete decommission; Fig. 1). For each reservoir, we calculated two impact scores: one for the reservoir creation and construction of the dam (ISR; where CFs were multiplied by the affected area) and one for the hydroelectricity production (IS; where ISRs were divided by the annual kWh produced for a given powerplant). We also took a multi-scale approach to examine if patterns observed at the sampling station, reservoir and biome scale were comparable.

**Figure 1.**
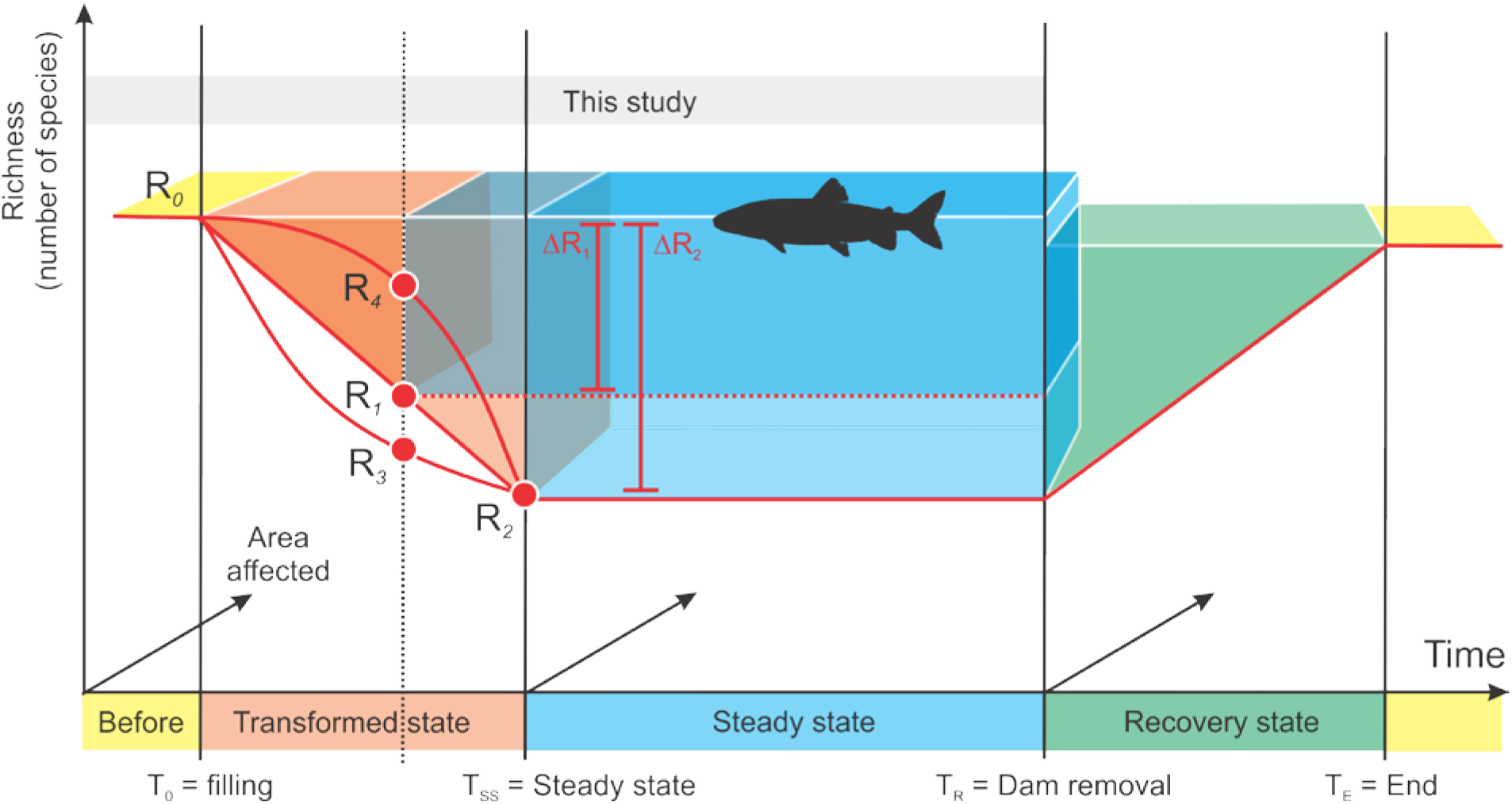
Schematic representation of an area-time framework representing the rate of change in richness experienced in a given reservoir. R_0_ represents the richness before impoundment, R_is_ represents different richness during the transformed state of the reservoir and where the fish community respond to environmental change following impoundment. The ΔQs represent the steady state where fish community should have reached a new equilibrium and where the rate of change in fish species should stabilize. The recovery state should start when the reservoir and dam will be decommissioned. This study addresses the period between the before impoundment to the reach of the steady state.

### 2.2 Richness data extraction and literature search

The literature search for this paper has been performed previously for another companion meta-analysis examining the global effect of dam on fish biodiversity (Turgeon et al., 2019b). In a nutshell, the search resulted in 668 publications (mostly peer-reviewed articles). For this paper, we excluded modelling and simulation exercises, and we refined our selection criteria to include only references that had unbiased quantitative data on the fish community before and after impoundment, and where the main purpose of the dam was to produce hydroelectricity. Data are limited to storage reservoirs technology and thus does not include run of the river and pumping station technologies due to a lack of longitudinal data. A total of 30 references met our selection criteria (Database A). See Turgeon et al. (2019b) for a detailed methodology about the literature search, and data extraction.

### 2.3 Extracting the area affected by the dam and reservoir

To extract the area affected by the construction of the dam and reservoir, we extracted the area occupied by the rivers and lakes prior to impoundment (hereafter called the affected area) both upstream and downstream of the dam (Fig. A). Change in land use from terrestrial to reservoir (inundated land area; ILA) is out of the scope of this paper, but see (Dorber et al., 2018) for a proposal to model net land occupation of hydropower reservoirs in LCA. To get the affected area information, we used various sources. For recent reservoirs, we used Google Earth Pro with the historical satellite imagery tool (Landsat/Copernicus images). Other sources of historical maps consisted in the USGS historical topographic maps for most of the United States reservoirs (https://viewer.nationalmap.gov/basic/), the Old Maps Online website for old reservoirs in Africa and South America (http://www.oldmapsonline.org/). The image of the river bed before impoundment was exported as a raster image in QGIS (v.2.18.16; http://www.qgis.org). The affected area was hand drawn as a polygon in a vector layer, and the total area of the polygon was extracted. Two polygons were extracted per reservoir, one upstream and one downstream of the dam. Upstream, we assumed that the impacts of the reservoir and the dam on fish community did not go beyond the impounded area and thus, used the upper end of the reservoir as the upper limit of the affected area. For downstream stations, we used 10 km downstream of the dam to set the lower limit of the polygon. We tested for the effect of different distances downstream of the dam (5, 10, 15, 25 km), in addition to the distance at which data were collected (mean ± SD; 13 km ± 45 km; median; 0.35 km) and they were all strongly correlated (Pearson r > 0.80; *unpublished analysis*).

### 2.4 Calculation of change in richness

#### 2.4.1 Sampling station scale

For each sampling station *i*, located either upstream or downstream of the dam in reservoir *j*, we calculated the rate of change in richness over time with a linear regression. The rate of change in richness was extracted using the estimated slope of the regression between richness and time (Equation 1) and we used the standard error of the estimate to calculate the 95% confidence interval (CI; see Database A). In this study, we assumed a linear relationship between richness and time, but some studies empirically observed a rise and fall of richness over time (Agostinho et al., 1994; Lima et al., 2016). See discussion for potential limitations and biased interpretation associated with this assumption.

#### 2.4.2 Reservoir and biome scales

When more than one station were sampled per reservoir, we used general linear mixed effect model (glmm; lmer function in lme4 package v.1.1-13; (Bates et al., 2018) to calculate the rate of change in richness over time, separately for upstream and downstream stations. At the reservoir scale, we controlled for temporal non-independence of the data by using sampling station as a random factor. At the biome scale, we controlled for spatial and temporal non-independence of the data by nesting each sampling station *i* into their respective reservoir *j*. All analyses were performed using R v. 3.3.2 (R Core Team, 2017).

### 2.5 Calculation of Characterization Factors (CF)

#### 2.5.1 Sampling station scale

To calculate CFs, we multiplied the observed rate of change in richness (ΔR/Δt; where ΔR stands the difference in richness and Δt stands for the duration of the study) by the time it take to reach a defined steady state *t*_*ss*_ (time horizon at which we considered that the rate of change in richness = 0; see Fig. 1) as per Equation 1, and divided the result by the average richness observed before impoundment for a given sampling station (R_0ij_). We did this for each sampling station *i* in reservoir *j*. The duration at which fish richness has been sampled for a given study (Δt) varied greatly across studies and biomes (*e.g.*, from less than five years to 40 years, see Database A). This can be problematic when comparing short duration studies with longer ones, because the longer the time after impoundment (Δt), the bigger the ΔR will be, which can result in an underestimation of PDFs (Fig. 1; see R_1_ vs. R_2_). To make studies comparable in their steady state, we tested with a sensitivity analysis, different scenarios of time to reach the steady state (*t*_*ss*_ = 5, 10, 20, 25 and 30 years after impoundment; Equation 1). To calculate the uncertainty associated with the CFs, we used the standard error (SE) from the estimate of the rate of change in richness (from the glmm) and multiplied it by the different scenario of time to reach the steady state and then divided it by the average richness observed before impoundment. From this scaled SE, we calculated the 95% CI.

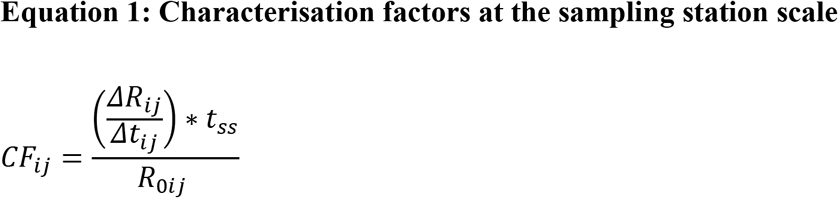

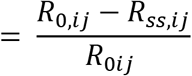

where (ΔR_*ij*_/Δt_*ij*_) is the observed rate of change in richness extracted in sampling station *i* in reservoir *j* by using the slope of the regression between the observed change in richness (ΔR) for a given period (Δt) and t_ss_ are the different steady state scenarios (5, 10, 20, 25 and 30 years after impoundment).

#### 2.5.2 Reservoir and biome scales

To test if the CFs are valid and robust across scales (sampling station, reservoir and biome), we also computed CFs at the reservoir and biomes scales. At the reservoir scale, we calculated a mean upstream CF, a mean downstream CF, as well as a mean CF (upstream CF + downstream CF when upstream and downstream stations were available). To do so, we averaged the CFs calculated for upstream sampling stations in reservoir *j*. We then squared the SE associated with the coefficient the regression (slope of the observed change in richness for a given period) for each upstream sampling station of reservoir *j* added them together to get the total variance for reservoir *j*. We then divided this variance by the number of sampling stations in reservoir *j* raised to the power of 2, and square rooted that variance to get the SE of the mean CF, and we calculated the 95% CI. We did the same procedure for downstream stations and for the biome scale. At the biome scale, we used CF_j_ as units (calculated at the reservoir scale) instead of CF_*i*_ (calculated at the sampling station scale).

### 2.6 Calculation of impact scores (ISR and IS)

We were also interested to evaluate the potential environment impact of creating a reservoir (ISR; elementary flow = area affected upstream and downstream of the dam) and of producing hydroelectricity (IS; elementary flow = kWh produced for a given reservoir). To do so, we multiplied the CF by the area affected by the reservoir and the dam (*i.e.*, area occupied by the rivers and lakes prior to impoundment, see Fig. A). Because we expect different impacts upstream and downstream of the dam, we calculated the impact score of creating and operating a reservoir (ISR; CF m^2^ yr) as the sum of downstream (ISR_*dj*_) and upstream impacts (ISR_*uj*_; Fig. A). Impact scores were calculated at the reservoir and biome scales.

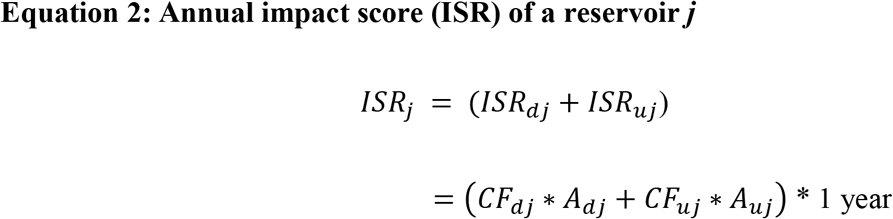

where *A* stands for the area affected upstream (*uj*) or downstream (*dj*) of the dam for reservoir *j* (Fig. A).

To calculate the impact score per unit of hydroelectricity produced by a powerplant associated to a given reservoir, the impact score ISR_j_ was divided by the annual electricity production, P*j* (kWh/year).

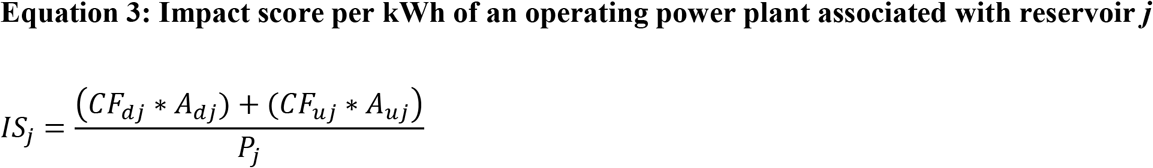

### 2.7 Sensitivity analysis of the Steady State Scenarios (SS)

Because we simulated CF and IS for different steady state scenarios (5, 10, 20, 25 and 30 y after impoundment) we evaluated the sensitivity of CF for each incremental increase in the time to steady-state (t_SS_).

## 3. Results

### 3.1 Rate of change in fish richness across scales and biomes

Upstream and downstream of the dam, the rate of change in fish richness over time varied strongly across sampling stations, reservoirs and biomes (Fig. 2). When all biomes, reservoirs and sampling stations were combined, richness significantly decreased over time at a rate of 0.29 species per year upstream of the dam (estimate ± SD = −0.293 ± 0.074, 95% CI = −0.439 to - 0.148) and at a comparable rate downstream of the dam (0.26 species per year; estimate ± SD = −0.264 ± 0.082, 95% CI = −0.424 to −0.104). In the boreal biome, there was no significant change in richness over time at all scales (sampling station, reservoirs and biome) and for both upstream and downstream stations (95% CI overlapped with zero; Fig. 2 a, b). In temperate and tropical regions, we observed a significant decrease in richness over time at the biome scale for upstream stations (loss of 0.26 and 1.6 species per year respectively; Fig. 2 c, e). Downstream of the dam, we observed a significant decrease in richness in the temperate region (loss of 0.34 species per year) but not in the tropics (Fig. 2 d, f). In these two biomes, some sampling stations and reservoirs showed either a significant decrease or, interestingly, an increase in richness over time following impoundment (Fig. 2 c-f).

**Figure 2.**
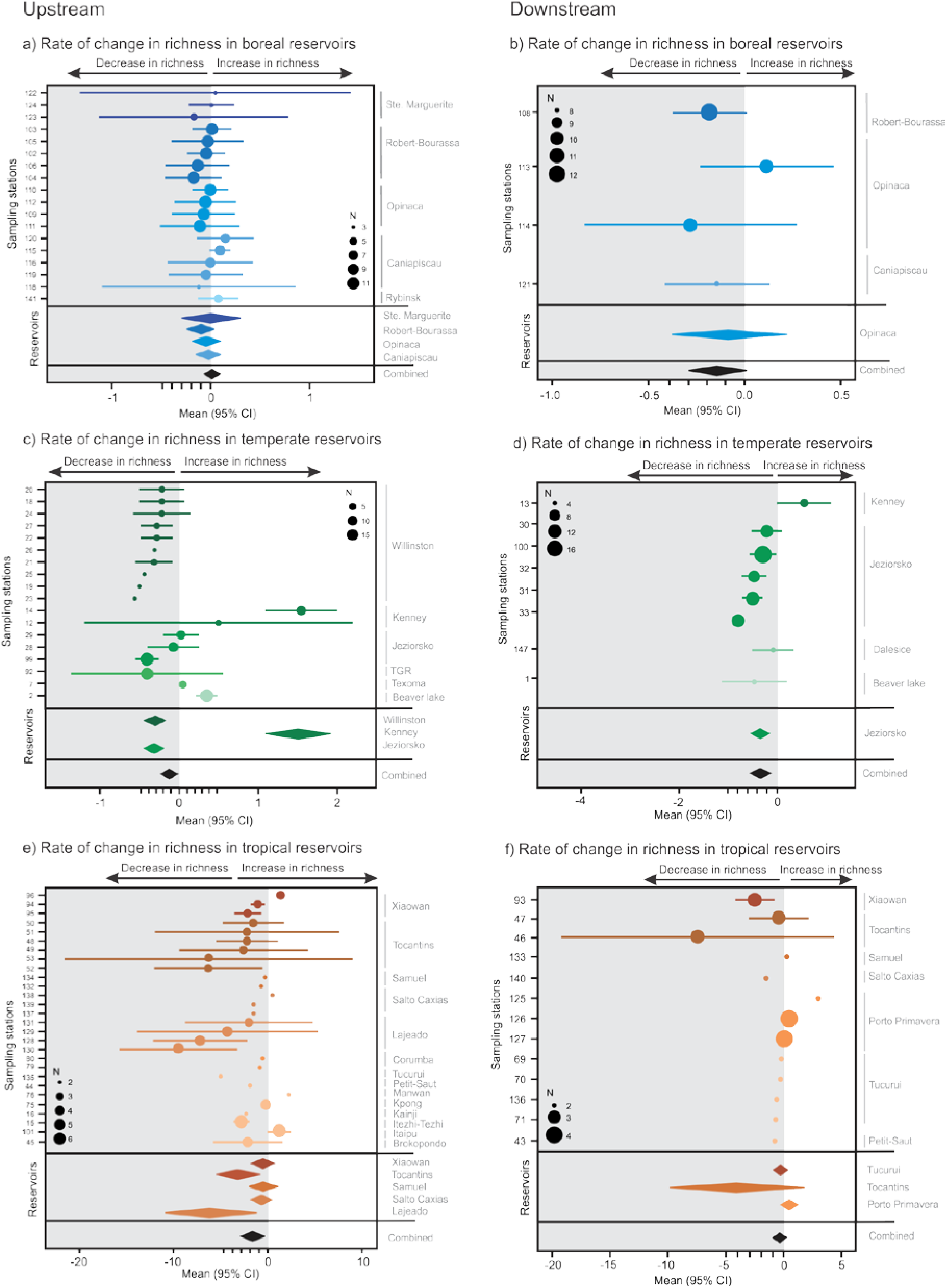
Empirically derived rates of change in richness and their 95%CI in upstream and downstream sampling stations, and reservoirs from three biomes: boreal (23 sampling stations, 5 reservoirs), temperate (26 sampling stations, 7 reservoirs) and tropical reservoirs (41 sampling stations, 15 reservoirs). The size of the circles represents the number of observations in the time series used to derive the rate of change in richness.

### 3.2 Characterization factors (CF)

The magnitude of the impact and statistical significance of CFs were sensitive to the assumption of reaching the steady-state (t_SS_), differed across biomes, but were consistent across scales and position (downstream or upstream of the dam; Fig. 3, Figs. B.1, B.2 and B.3). In boreal ecosystems, there was no significant loss of species upstream and downstream of the dam at the sampling station scale and for all steady state scenarios (Fig. B.1). When data were combined at the reservoir scale, no loss of species was observed upstream (Fig. 3 a), and a marginal loss of species was observed in one reservoir downstream of the dam (Fig. 3 b, Table 1). In temperate and tropical ecosystems, there were some significant gains and losses of species upstream (Fig. B.2a and Fig. B.3a) and downstream of the dam at the sampling station scale (Fig. B.2b and Fig. B.3b). When data were combined at the reservoir scale for temperate and tropical ecosystems, we also observed gains and losses of species (Fig. 3, Table 1).

**Figure 3.**
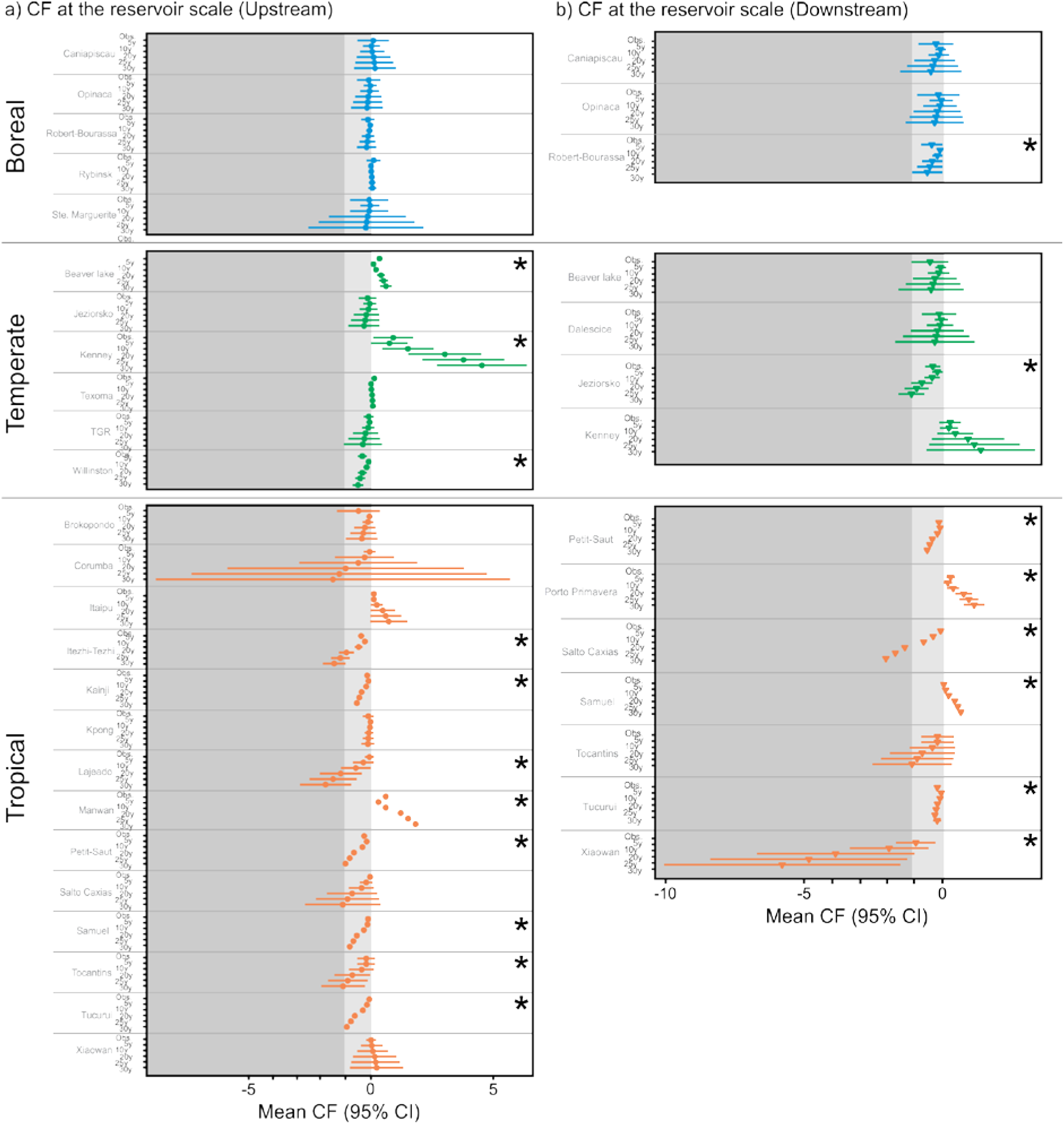
Characterization factor estimates (CFs) and their 95% CI at the reservoir scale for three biomes, a) upstream and b) downstream of the dam and for the observed duration of the study, and the 5 simulated steady state scenarios (5y, 10y, 20y, 25y and 30y). A negative value represents a loss of species and a positive value a gain in species. CF values in the dark grey area means that 100% of the species were loss. Stars beside the CF values indicate a statistically significant CF.

**Table 1.** Estimates ± Standard Error (SE) for Characterization factors (CF), Impact scores for the creation of the reservoir (ISR) and impact scores to produce hydroelectricity (IS) at the reservoir and biome scales.

| Reservoirs/biome | Biome | U, D, or<br>U+D | CF<br>(PDF*y)<br>Estimate ± SE | ISR<br>(PDF*km<sup>2</sup>*y) x<br>1.0E+08<br>Estimate ± SE | IS<br>(PDF*km<sup>2</sup>*y/<br>kwh)<br>Estimate ± SE |
| --- | --- | --- | --- | --- | --- |
| **BOREAL** | **B** | **U + D** | **-0.159 ± 0.204** | **-0.604 ± 12.30** | **0.061 ± 0.012** |
| Ste-Marguerite | B | U | -0.069 ± 0.401 | -0.240 ± 1.397 | -0.009 ± 0.051 |
| Rybinsk | B | U | 0.021 ± 0.027 | 0.890 ± 1.183 | 0.138 ± 0.184 |
| Robert Bourassa | B | U + D | -0.243 ± 0.247 | -3.360 ± 4.758 | -0.009 ± 0.013 |
| Opinaca | B | U + D | -0.148 ± 0.813 | -3.084 ± 15.77 | -0.008 ± 0.042 |
| Caniapiscou | B | U + D | -0.085 ± 0.850 | 4.630 ± 33.02 | 0.201 ± 1.436 |
| **TEMPERATE** | **T** | **U + D** | **0.524 ± 0.442** | **0.028 ± 0.029** | **0.102 ± 0.350** |
| Three Gorges | T | U | -0.109 ± 0.000 | -5.152 ± 0.000 | -0.005 ± 0.000 |
| Texoma | T | U | 0.026 ± 0.007 | 0.032 ± 0.008 | 0.013 ± 0.003 |
| Kenney | T | U + D | 1.974 ± 2.351 | 0.067 ± 0.042 | 0.559 ± 0.350 |
| Jeziorsko | T | U + D | -0.468 ± 0.575 | -0.048 ± 0.075 | -0.288 ± 0.440 |
| Dalesice | T | D | -0.093 ± 0.244 | 0.064 ± 0.068 | - |
| Beaver Lake | T | U + D | 0.068 ± 0.103 | 0.064 ± 0.018 | 0.035 ± 0.010 |
| **TROPICAL** | **TR** | **U + D** | **-0.781 ± 0.148** | **-2.620 ± 0.721** | **-0.056 ± 0.010** |
| Xiaowan | TR | U + D | -1.853 ± 0.987 | -0.024 ± 1.906 | 0.000 ± 0.010 |
| Tucurui | TR | U + D | -0.421 ± 0.164 | 10.35 ± 2.603 | -0.048 ± 0.012 |
| Tocantins | TR | U + D | -0.747 ± 1.011 | -2.210 ± 1.552 | -0.093 ± 0.065 |
| Samuel | TR | U + D | -0.069 ± 0.144 | -0.737 ± 0.194 | -0.081 ± 0.021 |
| Salto Caxias | TR | U + D | -1.065 ± 0.192 | -2.057 ± 0.403 | -0.038 ± 0.007 |
| Porto Primavera | TR | D | 0.381 ± 0.111 | 0.891 ± 0.259 | 0.008 ± 0.002 |
| Petit Saut | TR | U + D | -0.532 ± 0.174 | -0.350 ± 0.085 | -0.075 ± 0.018 |
| Manwan | TR | U | 0.608 ± 0.000 | 0.518 ± 0.000 | 0.007 ± 0.000 |
| Lajeado | TR | U | -0.619 ± 0.310 | -5.120 ± 2.604 | -0.116 ± 0.058 |
| Kpong | TR | U | -0.042 ± 0.046 | -0.008 ± 0.009 | -0.001 ± 0.001 |
| Kainji | TR | U | -0.192 ± 0.000 | -2.298 ± 0.000 | -0.165 ± 0.000 |
| Itezhi-Tezhi | TR | U | -0.501 ± 0.078 | -0.481 ± 0.076 | -0.076 ± 0.012 |
| Itaipu | TR | U | 0.243 ± 0.129 | 1.872 ± 0.996 | 0.002 ± 0.001 |
| Corumba | TR | U | -0.518 ± 0.000 | -0.390 ± 0.000 | -0.027 ± 0.000 |
| Brokopondo | TR | U | -0.124 ± 0.109 | -0.510 ± 0.451 | -0.057 ± 0.050 |

Sensitivity analysis suggested that simulated CFs for steady state scenario reached 15y after impoundment and beyond were unlikely because many reservoirs lost 100% of the original richness which was never been observed in any reservoirs (Fig. 3, Fig. C). Steady state scenario reached at 5y underestimated species loss when compared to the observed duration (Fig. 3, Fig. C). For these reasons, a steady state scenario reached at 10y will be used to calculate impact scores, and to compare the impact of impoundment across biomes and reservoirs.

### 3.3 Impact scores for the creation of the reservoir and for hydroelectricity production

Impact scores for the creation of reservoirs (ISR) and for hydroelectricity production (IS) differed across biomes and reservoirs (Fig. 4, Table 1). ISRs in boreal and temperate regions were not significant for the observed duration of the study (O; Fig. 4 a) and for the steady state scenario of 10y (SS10; Fig. 4 a). However, three tropical reservoirs showed a significant ISR when using the SS10 (Fig. 4 a). These results translated into an ISR of 0 for boreal and temperate biomes, and a significant ISR for the tropics (Fig. 4 a, ALL). The directionality and significance of IS were comparable to ISR for both the reservoirs and biome scales (Fig. 4 b, Table 1).

**Figure 4.**
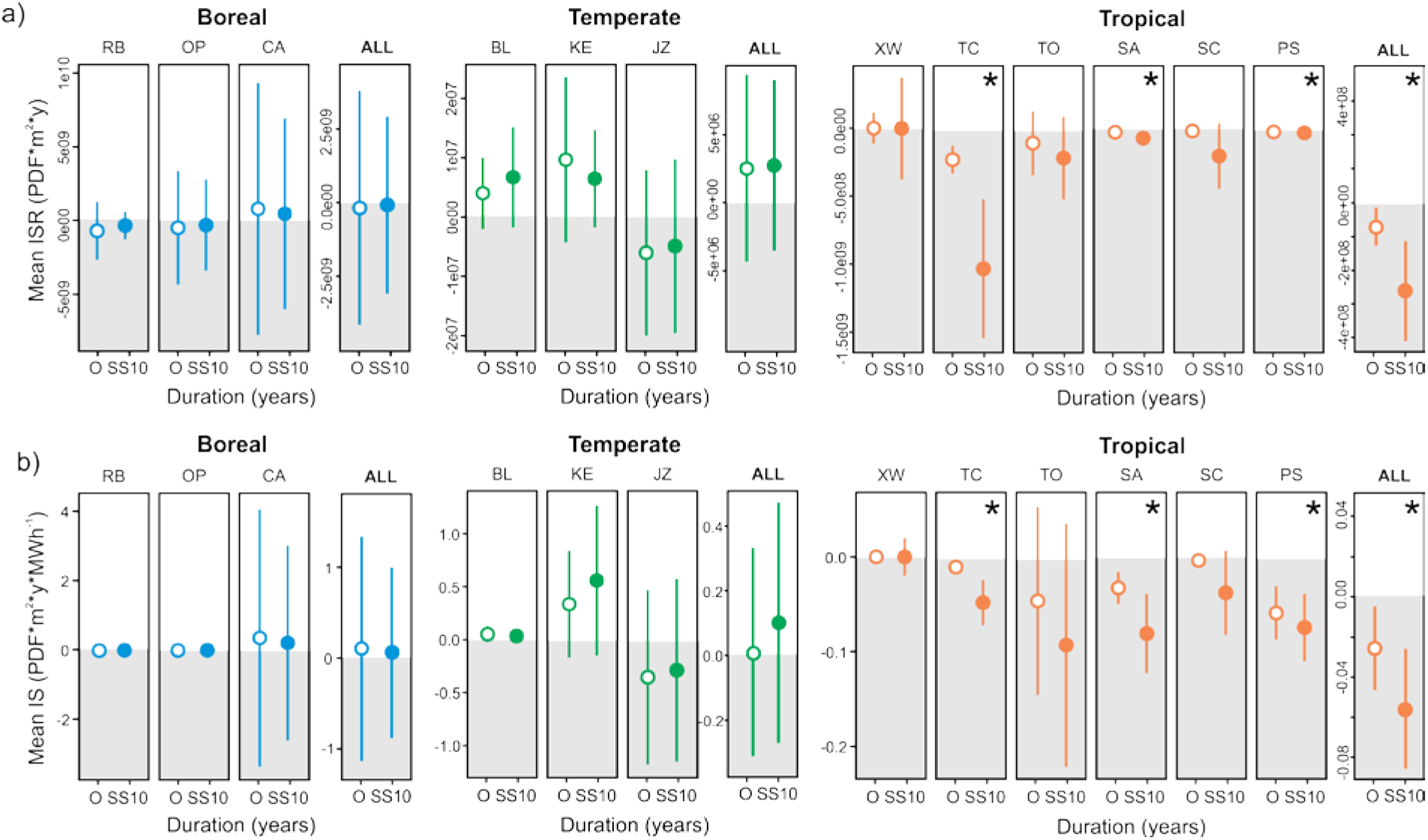
a) Mean reservoir and biome (ALL) impact score for the creation of the reservoir (ISR) and b) mean reservoir and biome (ALL) impact score of hydroelectricity production for the observed duration of the study and for the steady state scenario of 10 y in the three biomes. RB = Robert-Bourassa, OP = Opinaca, CA = Caniapiscau, BL = Beaver Lake, KE = Kenney, JZ = Jeziorsko, XW = Xiaowan, TC = Tucurui, TO = Tocantins, SA = Samuel, SC = Salto Caxias, PS = Petit-Saut. Stars beside the CF values indicate a statistically significant CF.

## 4. Discussion

### 4.1 Regionalisation is needed

Based on available empirical data (89 sampling stations located upstream and downstream of the dam and belonging to 27 reservoirs across three large biomes), we demonstrated that regionalization is needed for this impact category in LCA because the observed rate of change in fish richness in hydroelectric reservoirs varied across biomes, being minimal in boreal, marginal in temperate ecosystems, and significant in the tropics. This result suggests that hydroelectricity production in Northern countries located in the boreal biomes (*e.g.*, Canada, Russia, Norway, Sweden, Finland, Iceland), which account for more than 15% of the installed hydroelectricity production capacity in 2016 (International Energy Agency (IEA), 2016), has limited impacts on fish biodiversity. On the other hand, our dataset demonstrated that hydroelectricity production in the tropics has significant impacts on fish biodiversity at all scales. Rivers in species-rich tropical region located in Brazil (installed capacity of 91.7 GW, 85% of the generated energy in Brazil, 8% globally) and China (installed capacity of 319 GW, 17% of the generated energy in China, 27% globally), have been extensively harnessed for hydroelectricity production (Stickler et al., 2013; Winemiller et al., 2016; Ziv et al., 2012). Future hydroelectric development (planned and currently in construction) is concentrated in China, the Mekong region, Latin America and Africa, and the largest potential for future development is in Asia (International Energy Agency (IEA), 2016). All these regions have high fish richness and endemic species, some of these regions are recognized as biodiversity hotspots, and they will be particularly impacted by climate change regarding loss in water availability (Xenopoulos and Lodge, 2006). In a collective effort to decarbonize the worldwide economy and reduce GHG emissions, we urgently need appropriate supporting decision tools that consider long term economic, environmental and social costs (Fearnside, 2016; Kahn et al., 2014). The use of our developed CFs and ISs in LCA, accounting for potential impact of hydropower on aquatic ecosystems biodiversity, could help in this respect.

### 4.2 First empirically derived CF and IS

Apart from few unpublished attempts (Humbert and Maendly, 2008), this contribution is the first to empirically address the impact of hydroelectricity production on biodiversity in LCA. Recent methods and contributions in LCA addressed the impact of water shortages or consumption on biodiversity using Species-Discharge relationships (SDR; (Hanafiah et al., 2011; Tendall et al., 2014)) or Species-Area relationships (SAR; (de Baan et al., 2013; Verones et al., 2013)) but none of these contributions addressed the time it takes to reach the steady-state (Souza et al., 2015). It is also quite risky to relate potential change in water discharge to change in species richness using SDR because these curves reflect evolutionary and ecological outcomes roughly in equilibrium with natural discharge (Rosenberg et al., 2000; Xenopoulos and Lodge, 2006). Data limitations to build SDR curves are severe, especially for change in biodiversity. Species richness numbers are not readily available for most rivers of the world, and temporal sequences spanning changes in discharge are extremely rare. Data limitations thus make difficult any rigorous tests of species–discharge models (Xenopoulos and Lodge, 2006). Moreover, we still do not know the impact pathways and the main drivers of potential changes in biodiversity in reservoirs and regulated rivers. The impacts of damming a river go well beyond changes in water discharge. Dams and reservoirs drastically change the hydrological regime and the riverscape connectivity and may change the strength of trophic interactions upstream and downstream of the dam (Gracey and Verones, 2016; Renöfalt et al., 2010; Turgeon et al., 2019b, 2019a). These alterations may be much more important than change in discharge in affecting change in richness. Unless the impact pathway is convincingly understood, or SDR strongly validated with empirical data, we must be extremely careful in our choice of fate and effects factors in LCA.

### 4.3 Importance of temporal and spatial scaling in LCA

Great insights are achieved when multiple spatial and temporal scales are considered and/or compared because patterns observed at one scale are often not transferable to another scale (upscaling, downscaling issues; (Levin, 1992). In this study, the calculation of the CFs and ISs was strongly sensitive to the duration of the study but not to the spatial scale examined (*i.e.*, sampling station, reservoir and biome). We assumed a linear rate of change in richness over time since impoundment. This assumption would not be problematic if the duration of the study was long enough to convincingly reach the steady state phase (*i.e.*, new species assemblage equilibrium where the change in richness stabilizes after impoundment; Fig. 1) or if the duration of study was comparable across studies. However, the observed duration of the studies varied greatly (from only one year after impoundment, to 54 years after impoundment; Database S1) and the steady state was likely not reached in many reservoirs, especially in the tropics. This imply that CFs and ISs developed from short duration study will be underestimated (see Fig. 1; R_1_ vs. R_2_ resulting in two ΔQs). This pattern will be exacerbated if the relationship is non-linear (sigmoid, a rise and fall, or a non-linear accelerating decreasing rate; Fig. 1; R_4_ vs. R_2_) which is highly plausible (Agostinho et al., 1994; Lima et al., 2016). Most of the time series do not allow to test for non-linearity because they were too short, or the time steps between sampling events were too long. We also do not have the data to test if the time it takes to reach the steady state is similar across latitudes (*e.g.*, might be faster in the tropics and slower in boreal regions). To circumvent these problems, and to compare CFs and ISs across studies, we tested the sensitivity of different steady-state scenarios (5, 10, 20, 25 and 30y after impoundment) and assumed that using 10y after impoundment for all studies was a plausible scenario. We demonstrated that the impacts changed in magnitude depending on the duration of the studies and a standardization must be considered in LCA.

Some patterns observed in upstream stations were not corroborated by patterns observed in downstream stations suggesting that potentially different impact pathways affect the fish community upstream and downstream of the dam. The impacts upstream of the dams might be more closely related to the transformation of a lotic (river characteristics) into a lentic (lake characteristics) environment and to water levels fluctuations, whereas downstream impacts might be related to variation in water discharge (hydropeaking or not), and the dam acting as a barrier to fish migration/movement. In this study, we assumed that the extent of the impacts of damming the river was limited to the reservoir (upstream of the dam) or to 10 km downstream of the dam. We have very limited information on the extent to which the impacts of impoundment can be detected on fish community. Some studies detected significant changes in fish community and richness after impoundment upstream of the reservoir (Araújo et al., 2013; Lima et al., 2016; Penczak and Kruk, 2005) and as far as 25 km downstream of the dam (de Mérona et al., 2005). However, the impacts on fish community upstream of the reservoirs and downstream of the dam is probably site-specific because they will depend on how the dam is managed (*e.g.*, hydropeaking or not) the and the connectivity to tributaries. More studies are needed to determine the spatial extent, the impact pathways, and the factors contributing to changes in fish community when damming a river, upstream and downstream of the dam.

In this study, the observed empirical changes in richness from 89 sampling stations (upstream and downstream of the dam) were transferable to the reservoirs studied and were also transferable, but to a lesser extent, to the biomes. Our spatial coverage is thus global but the resolution (grain) of the CFs and ISs was coarse given the limited amount of empirical data. As empirical data and evidence will accumulate, the next step would be to refine the resolution at the scale of major habitat types (MHTs) or freshwater ecoregions of the world (FEOW; Abell et al., 2008) and to consider other taxa (macroinvertebrates, aquatic and riparian vegetation).

### 4.4 Limitations of developed CFs and ISs

Even though experts agreed on using species richness as a good starting point to model biodiversity loss in LCA (Teixeira et al., 2016), the use of Potentially Disappeared Fraction of species (PDF) is problematic for several reasons. First, it is imprudent to interpret a pattern of increased species richness (or no change in richness) as an indication of no impact of hydroelectricity production on biodiversity, if the pattern results from an increase in non-native species (*i.e.*, not from the initial regional pool of species, including exotic). We used change in fish richness but did not discriminate between native and non-native species because this information was not provided for all studies. In boreal reservoirs, no non-native species have been observed so the developed CFs and ISs are considered robust (Tereshchenko and Strel’nikov, 1997; Turgeon et al., 2019a). In temperate reservoirs, the observed increase in richness after impoundment in Beaver lake, Kenney and Texoma reservoirs (Figs 2, 3 and 4), was actually due to an increase in non-native species (Gido et al., 2000; Martinez et al., 1994; Rainwater and Houser, 1982). In tropical reservoirs, an increase in non-native species have also been observed in Itaipu, Manwan and Xiaowan reservoirs, all showing an increase in richness over time (Li et al., 2013; Lima et al., 2016; Xiaoyan et al., 2010). A companion study (Turgeon et al., 2019b), looking at a larger dataset and including reservoirs used for other purposes (*e.g.*, irrigation, flood control), found a gradient of impact on fish biodiversity from being minimal in boreal, intermediate in temperate and important in tropical reservoirs. The small CFs and ISs in temperate reservoirs may be underestimated and should thus be interpreted with caution. Future studies should look at the fate of both native species and non-native species to develop the CFs and ISs.

Second, looking at PDF do not account for potential change in fish assemblages (potentially affected fraction of species; PAF) or in species that are more vulnerable (endemic and/or threatened). Several alternatives indices and models have been suggested and used to account for loss in biodiversity in LCA (*e.g.*, functional diversity, ecosystem scarcity) (Souza et al., 2015) but data requirement is tremendous, species have different adaptive capacity in different regions of the world and will respond to impoundment differently, and most importantly we must deal with the incommensurable challenge of developing CFs and ISs locally or regionally but apply them globally with the same rigor and criteria.

Finally, our developed CF and ISs evaluated the impacts of hydroelectricity production in storage reservoirs and only on the aquatic ecosystem’s biodiversity (affected area; river and lakes transformed into reservoirs) and not on the terrestrial area transformed into a reservoir. A simplistic assumption could consider a loss of 100% of the impounded terrestrial habitat and a gain of 100% aquatic habitat. The biodiversity impact on the flooded area is very relevant issue, and some promising work have been done in this respect to model net land occupation of reservoir in Norway (Dorber et al., 2018).

## 5. Conclusions

By using empirical data on the rate of change in fish richness over time, with data before and after impoundment, on more than 89 sampling stations located upstream and downstream of the dam, and belonging to 26 reservoirs across three large biomes, this study is the first to propose robust and empirically developed characterization factors and impact scores of the effects of hydroelectricity production on aquatic biodiversity. Our results suggest that the impact of hydroelectricity production on fish richness is significant in tropical reservoirs, marginal in temperate and not significant in boreal reservoirs which calls for regionalization in LCA. Our results also demonstrated that the calculation of PDFs, and consequently ISs, was sensitive to the time it takes to reach the steady state for fish communities. A steady state scenario of 10 years after impoundment was the most plausible scenario based on the examination of PDFs at the sampling station and reservoir scales. Finally, PDFs and ISs were relatively robust to upscaling and downscaling issues (*i.e.*, patterns were consistent in their directionality across sampling stations, reservoirs and biomes), but the statistical significance of the impact changed across scales. Hydropower can be part of the solution to decarbonize our global economy but will come at substantially higher ecological cost to the tropics (Pelicice et al., 2017; Winemiller et al., 2016; Ziv et al., 2012).

