## Supplemental material for "Empirical characterization factors assessing the effects of hydroelectricity on fish richness across three large biomes"

^1^Institut de recherche sur la Forêt Tempérée - Université du Québec en Outaouais, 85 Rue Principale, Ripon, QC, CANADA, J0V 1V0, ^2^Hydro-Québec, Environment and Corporate Affairs, 75 René-Lévesque, Montréal, QC, CANADA, H2Z 1A4, ^3^CIRAIG, 3333 Chemin Queen-Mary, Montréal, QC, CANADA, H3V 1A2, ^4^Mathematical and Industrial Engineering Department, Polytechnique Montreal, 2900, boul. Édouard-Montpetit, 2500, chemin de Polytechnique, Montréal, QC, CANADA, H3T 1J4, ^5^UQAM, École des sciences de la gestion, 315, rue Sainte-Catherine Est, Montréal, QC, CANADA, H2X 3X2

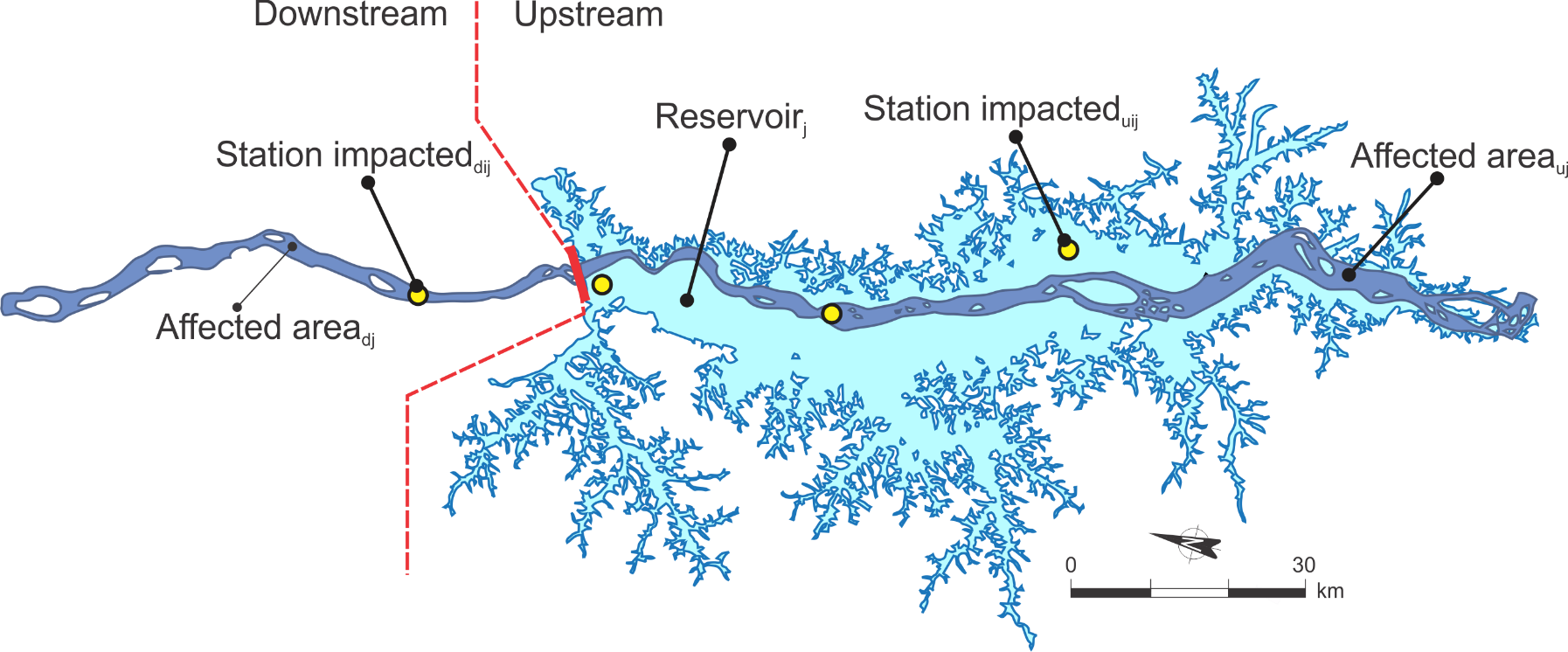

**Figure A**. Schematic representation of the affected area upstream and downstream of the dam, using the Tucurui reservoir in Brazil as an example. The reservoir polygon (light blue; after impoundment) has been extracted from the GRAND database (Lehner et al. 2010). The aquatic ecosystem before impoundment polygon (dark blue) has been hand drawn from an historical map (this study). CFs have been developed for each sampling stations (yellow circles) of a given reservoir, and then average to get CF at the reservoir scale.

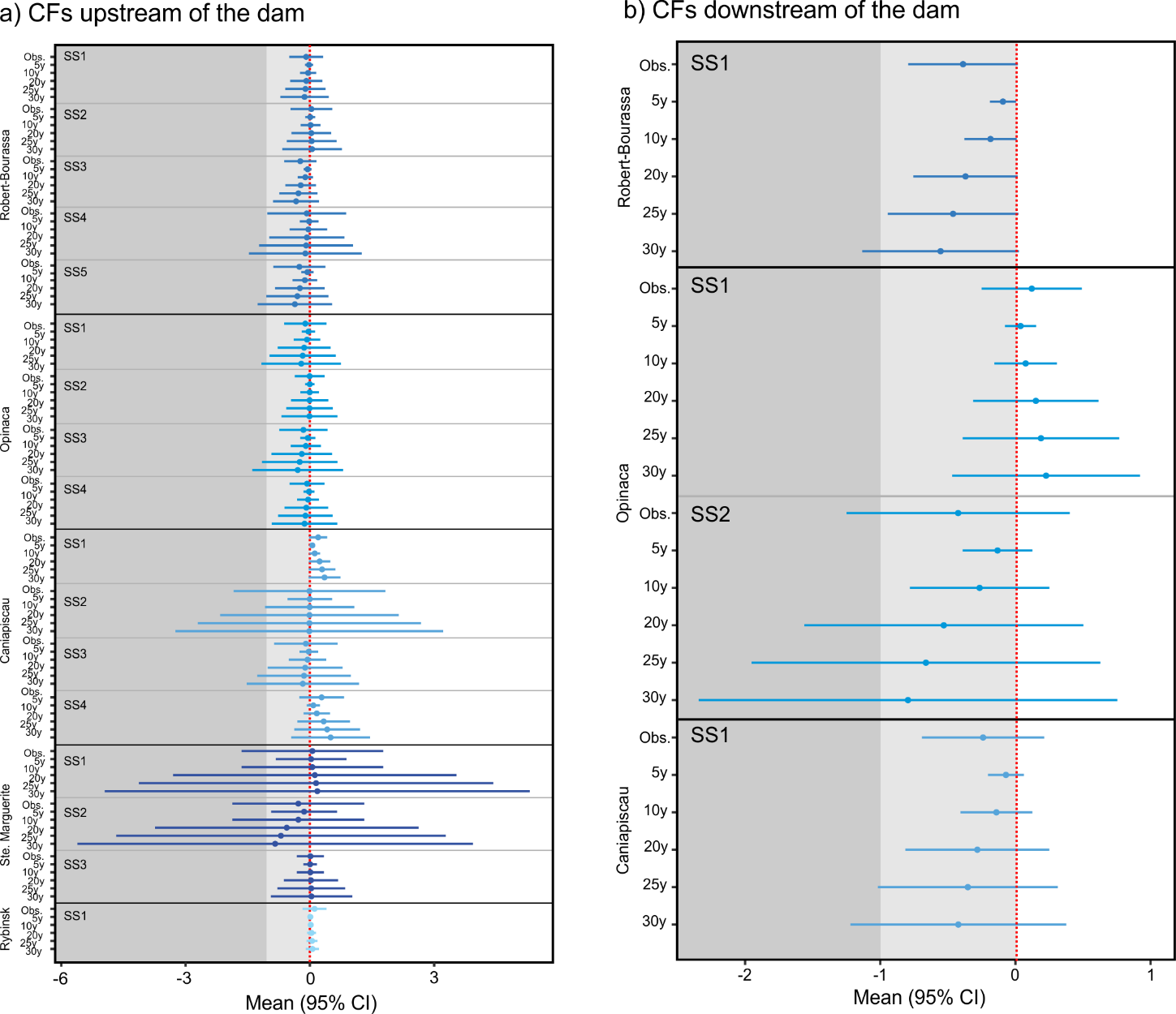

**Figure B.1. Characterization factors using the potentially disappeared fraction of the species (PDF) in the boreal region at the sampling station scale for a) upstream, b) downstream stations.** The observed duration (Obs.) and five steady state scenarios are presented (duration of 5y, 10y, 20y, 25y and 30y). The light grey area represents a loss of species and the dark grey area represent a loss of 100% of the species.

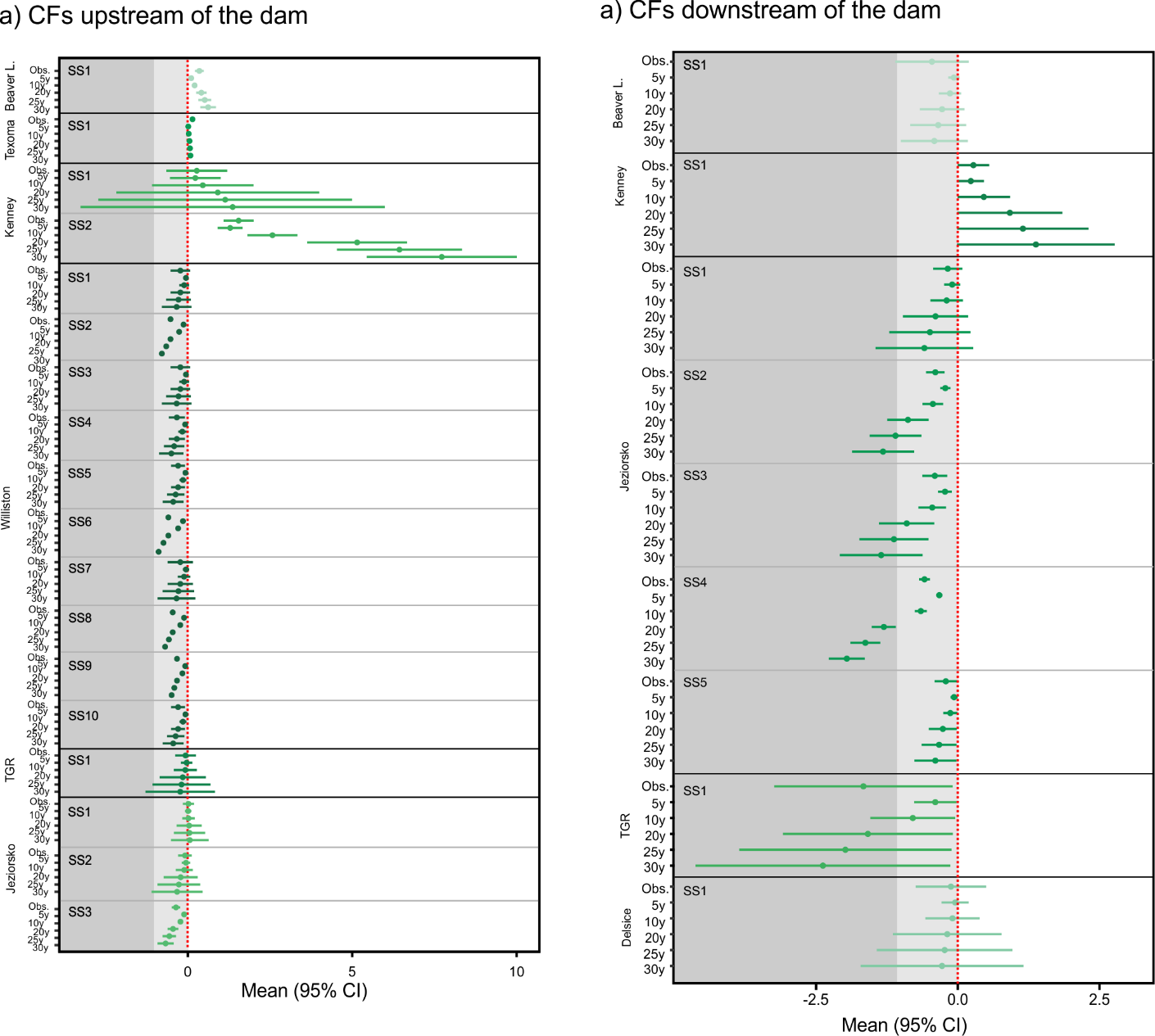

**Figure B.2. Characterization factors using the potentially disappeared fraction of the species (PDF) in the temperate region at the sampling station scale for a) upstream, b) downstream stations.** The observed duration (Obs.) and five steady state scenarios are presented (duration of 5y, 10y, 20y, 25y and 30y). The light grey area represents a loss of species and the dark grey area represent a loss of 100% of the species.

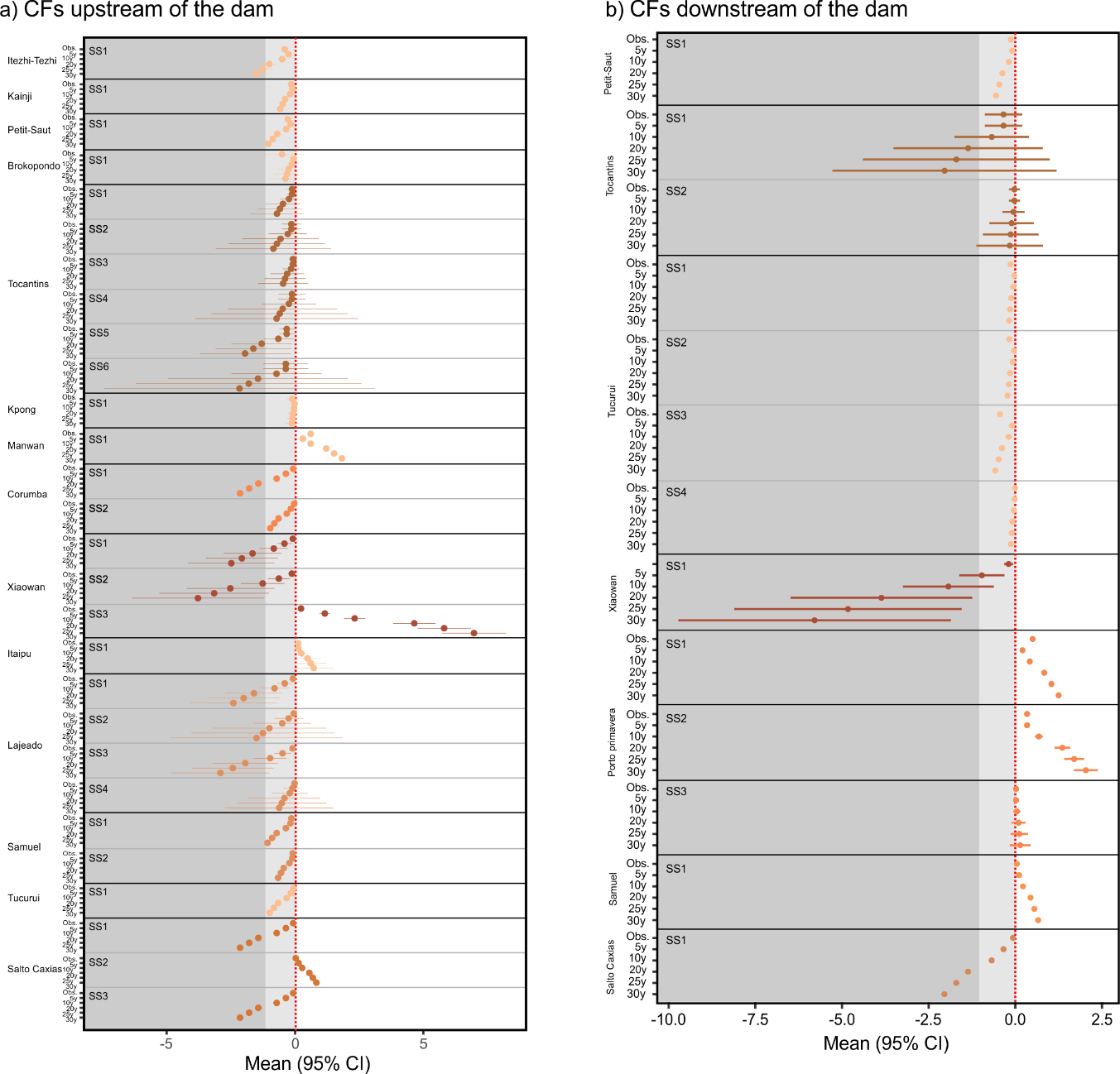

**Figure B.3. Characterization factors using the potentially disappeared fraction of the species (PDF) in the tropical region at the sampling station scale for a) upstream, b) downstream stations.** The observed duration (Obs.) and five steady state scenarios are presented (duration of 5y, 10y, 20y, 25y and 30y). The light grey area represents a loss of species and the dark grey area represent a loss of 100% of the species.

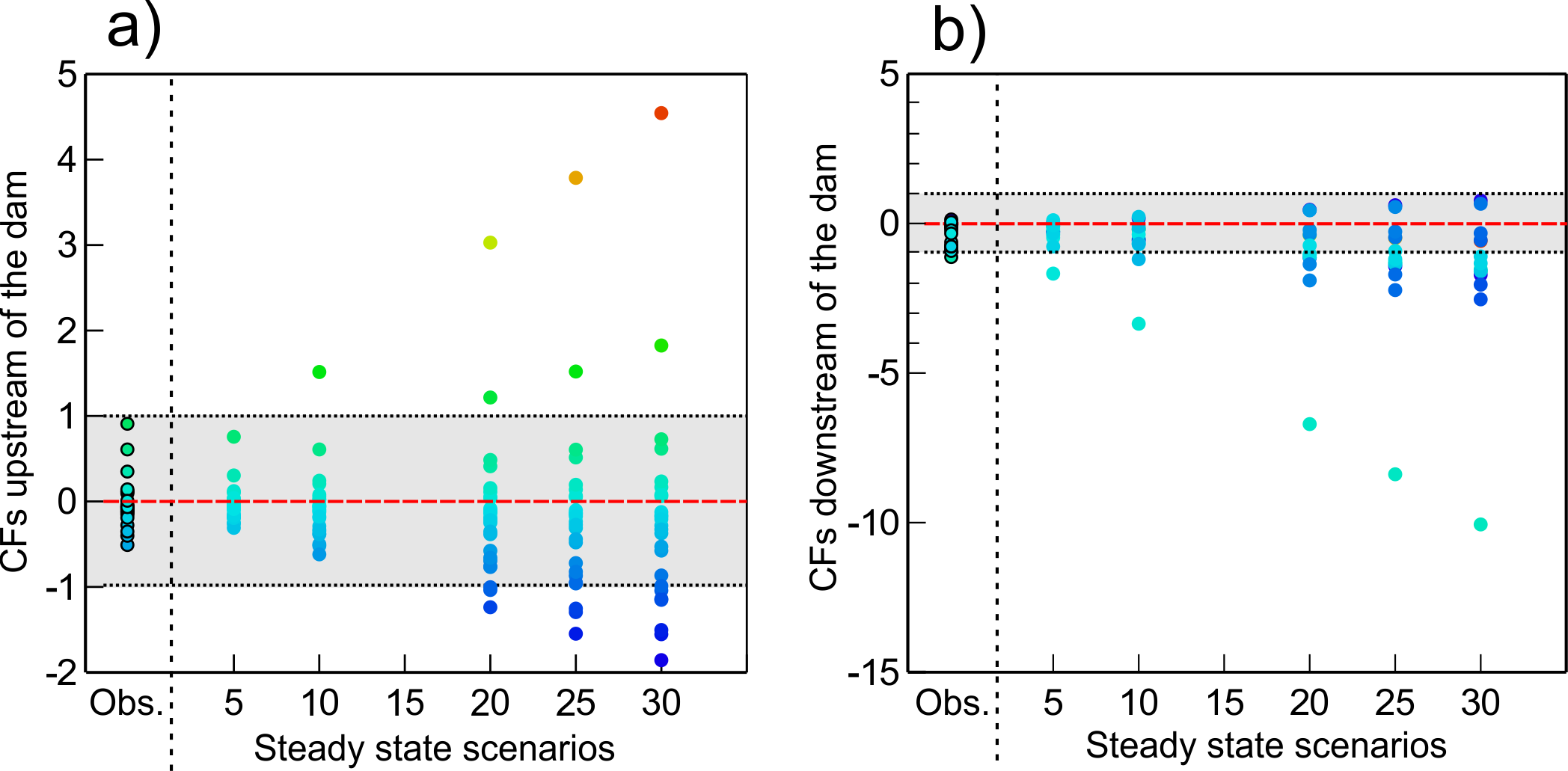

**Figure C.** Sensitivity analysis examining the effects of different steady state scenarios on the calculation of Characterization factors (CFs) for a) upstream stations and b) downstream stations.
